# *Burly1* is a mouse QTL for lean body mass that maps to a 0.8-Mb region on chromosome 2

**DOI:** 10.1101/231647

**Authors:** Cailu Lin, Brad D. Fesi, Michael Marquis, Natalia P. Bosak, Anna Lysenko, Mohammed Amin Koshnevisan, Fujiko F. Duke, Maria L. Theodorides, Theodore M. Nelson, Amanda H. McDaniel, Mauricio Avigdor, Charles J. Arayata, Lauren Shaw, Alexander A. Bachmanov, Danielle R. Reed

## Abstract

Our goal was to fine map a mouse QTL for lean body mass (*Burly1*) using information from several populations including newly created congenic mice derived from the B6 (host) and 129 (donor) strains. The results from each mapping population were concordant and showed that *Burly1* is likely a single QTL in a 0.8-Mb region at 151.9-152.7 Mb (*rs33197365* to *rs3700604*) on mouse chromosome 2. Results from mice of all the mapping populations we studied including intercrossed, backcrossed, consomic, and congenic strains indicate that lean body mass was increased by the B6-derived allele relative to the 129-derived allele. We determined that the congenic region harboring *Burly1* contains 26 protein-coding genes, 11 noncoding RNA elements (e.g., lncRNA), and 4 pseudogenes, with 1949 predicted functional variants. The effect of the *Burly1* locus on lean body weight was apparent at all ages measured and did not affect food intake or locomotor activity. However, congenic mice with the B6-allele produced more heat per kilogram of lean body weight than did controls, pointing to a genotype effect on lean mass metabolism. These results show the value of integrating information from several mapping populations to refine the map location of body composition QTLs.

## Introduction

An average adult mouse weighs about 25-30 g, and most of that weight is lean body mass. Lean body mass can differ almost 3-fold among common inbred strains, and this is highly heritable [1, 2]. By interbreeding specific strains, it has been identified dozens of influential genomic regions, or quantitative trait loci (QTLs), many of which are cataloged in the Mouse Genome Database [3]. Identifying the underlying causal genetic variants responsible for lean body mass [4], while challenging, is an important scientific goal, because lean body mass affects many tissues and functions of the body that, in turn, affect basal metabolic rate, metabolic health [5, 6], the immune system, and bone development [7].

Several genes and their variants greatly affect body size or lean body mass composition. Perhaps the best known are alleles of the myostatin gene that markedly increase muscle mass in mice [8, 9], cows [10-12], sheep [13], and other animals [14], including humans [15]. Other well-known variants are components of the growth hormone pathway, such as the *little* mutation [16, 17] and *dwarf* [18]. ‘*Little*’ mice are small with reduced lean body mass whereas *dwarf* mice are tiny [19] but have the usual proportion of lean and fat mass [20]. Beyond these single-gene mouse mutations, natural variation in lean body mass has a complex genetic architecture with numerous genes involved. Exactly how many genes contribute is unclear: QTL experiments suggest scores of loci [21], and knockout experiments suggest that almost a third of viable strains have reduced body weight or composition [22, 23], indicating that many thousands of genes may participate. Investigators using meta-analysis of human genome-wide association approaches indicate there are currently five reproducible loci for human lean body mass [24] but even collectively these studies are underpowered in part because direct measures of lean body mass are time-consuming and require specialized equipment relative to other measures like height. We speculate that when there are a comparable number of subjects studied, human lean mass will be similar in genetic architecture to human height [25, 26].

In addition to genotype, other factors affect the amount of lean body mass of individual mice, including sex, age, and maternal characteristics. Investigators consistently observe sex effects, with male mice having more lean body mass than females of the same strain [1]. There are also consistent age effects, with lean body mass peaking in middle to late adulthood and then declining [27]. Maternal effects are potent too, and the dam’s age, diet, behavior, and litter size account for some variation in the amount of lean body mass of her offspring [28]. Another factor that affects body composition is the parental origin of particular inherited alleles (imprinting) [29, 30] with paternally transcribed alleles generally favoring rapidly growth.

Genotype interacts with these factors, such as sex and age, to affect lean body composition [31]. Therefore, to better isolate genetic effects we keep these factors as constant as we can. Thus, in the current work, we compare 180 days old male littermates segregating a particular genetic variation so that age, sex and maternal effects are similar but the genotype is different. However, it is impractical to compare littermates differing in genotype of some specialized mouse strains, such as consomics, so in these cases we compare homozygous consomic mice to inbred mice of the host strain (e.g., [32, 33]). This practice has limitations but is sometimes expedient.

With these points in mind, our goals here were to map a particular lean body mass QTL (*Burly1*) and to identify the underlying genetic variants using a pair of contrasting inbred mouse strains. We began by intercrossing the heavier C57BL/6ByJ (B6) strain with the lighter 129P3/J (129) strain [34-36] and found a QTL on chromosome 2 for body weight, *Bwq5*. Mouse chromosome 2 harbors many related QTLs (e.g., [37-40]). This density makes it especially hard to dissect particular genetic effects on body composition and thus this intercross population did not provide sufficient mapping resolution to narrow the genomic interval. Also the phenotype, body weight as a proxy measure of lean body mass, was imprecise. Therefore, our next steps were to dissect this QTL by creating and studying additional mapping resources and by using direct measures of body composition rather than body weight. To that end, we studied several additional strains: a second intercross population, two reciprocal consomic [41], and several congenic strains, as well as the backcross mice produced during the breeding of these strains.

To measure lean body mass, we used both dual-energy X-ray absorptiometry (DEXA) and magnetic resonance (MR). To choose these methods, we considered the four common ways to measure lean body mass in rodents: total body weight, DEXA [42], MR [43, 44], and chemical extraction [45]. As mentioned above, total body weight is an imperfect proxy measure of lean body mass because it includes adipose tissue (fat mass), which can differ among rodents strains up to 20-fold [1, 2]. Most investigators consider chemical extraction the gold standard [42, 46] but it has at least two limitations: it can be conducted on dead mice only and it is time-consuming [45]. DEXA and MR are more direct measures than body weight and are comparable in validity compared with chemical extraction methods and can be conducted with living mice. DEXA requires mice to be anesthetized whereas MR can be used with mice that are awake. Thus, MR requires less time and fewer resources than does DEXA.

MR and DEXA provide estimates of total lean mass. To get a more complete picture of the *Burly1* phenotype, it is useful to measure the mass of individual organs. Many organs contribute to lean body mass, and necropsy results have the potential to identify overgrowth of a particular organ (i.e., allometry). It is also helpful to measure food intake and locomotor activity, because energy intake and expenditure is linked to learn body mass. Likewise information about the digestion and oxidation of energy sources are useful clues to the underlying genetic cause of differences in lean body mass [47].

Here we report all the *Burly1* mapping data we obtained from 2,070 mice derived from 2 intercrosses, 4 backcross generations, one consomic and one sub-consomic strain (defined below), and 25 congenic strains. To compare these mapping populations, we assayed or imputed genotypes for a common set of markers on mouse chromosome 2, and analyzed the genotype-phenotype associations using several approaches. Where possible, we compared littermates by genotype and took other steps to reduce sources of variation arising from sources other than inborn genotype. To gain insight about how the *Burly1* locus affects lean body mass, we characterized its effects in mice at different ages, and measured related traits.

## Methods

### Animal Husbandry

We bred all mice in a vivarium at the Monell Chemical Senses Center, located in Philadelphia, Pennsylvania (USA), using inbred B6 and 129 mice originally purchased from the Jackson Laboratory. Husbandry practices were stable throughout the study, with nearly the same vivarium personnel, cages, and type of bedding (Aspen Shavings, Northeastern Products Corp, Warrensburg, NY). All mice were fed Rodent Diet 8604 (Harlan Teklad, Madison, WI) and lived in a 12:12 light cycle, with lights off at 7 pm (barring unusual circumstances, e.g., power outages). For most mapping populations, we studied body composition of male mice only, to reduce overall trait variation and increase mapping power. However, in some experiments we collected data from female mice for other reasons, and we report those data here as well. The Monell Institutional Animal Care and Use Committee approved these study procedures.

### Intercrosses

We reported the body weight results from the first F2 intercross [34, 35, 48] but we also include them here to compare with results from later mapping populations. We bred a second intercross population with the same parental strains and measured lean and fat mass using DEXA. For this second intercross, we previously reported only the statistical results for fat but not lean body mass [49].

### Backcrosses and Consomics

To make consomic strains, we produced reciprocal N_2_ (F_1_ x B6 and F_1_ × 129) and then N_3_ backcross generations, followed by serial backcrossing and intercrossing of male and female mice to create the consomic B6.129-Chr2 and 129.B6-Chr2 strains. We were successful at creating only of the two reciprocal consomic strains (B6.129-Chr2) which is publically available and listed in **S1 Table** [50]. The unsuccessful strain was the129.B6-Chr2 strain. It was difficult to breed so instead we created a strain with a partial rather than full-length donor chromosome which we refer to as ‘sub-consomic’. We measured lean body mass in male and female mice from both the fully consomic and the sub-consomic strain, inbred host strains or littermate controls, as well as male mice from the backcross generations used to create them (i.e., the N_2_ and N_3_ generations).

### Congenics

We bred congenic mice by backcrossing N_8_F_2_ males (heterozygous males with a partial donor chromosome 2 generated from the consomic strain B6.129-Chr2) to the B6 background to obtain 129-derived donor regions of various lengths. The goal was to identify male breeders with a donor region that overlapped the QTL location from the first F_2_ intercross. We named the strains with codes that reflect their lineage; for instance, all strains with the prefix 1 (e.g., 1.1) descended from common progenitors. We bred all congenic mice from B6 (inbred) mothers and from fathers that were heterozygous for the 129-derived donor region. This approach reduced maternal effects (all mothers of the congenic mice were the same genotype) and reduced imprinting variation (only fathers contributed the congenic donor region). This strategy allowed us to compare littermates with one copy of the donor region (heterozygous; 129/B6) to those without the donor region (homozygous; B6/B6; **Figure 1**). Each congenic mouse potentially was genetically unique (because the paternal donor region could shorten due to meiotic recombination). Therefore, we genotyped each congenic mouse to ensure we could define the donor region breakpoints. In addition to these congenic strains, we bred homozygous mice from a congenic strain with a small donor fragment. This strain is also now publicly available (strain 2.5; **S1 Table**). We produced additional mice from the littermates of strain 2.5 without the 129-derived donor fragment, for use as a comparison group.

**Figure 1.**
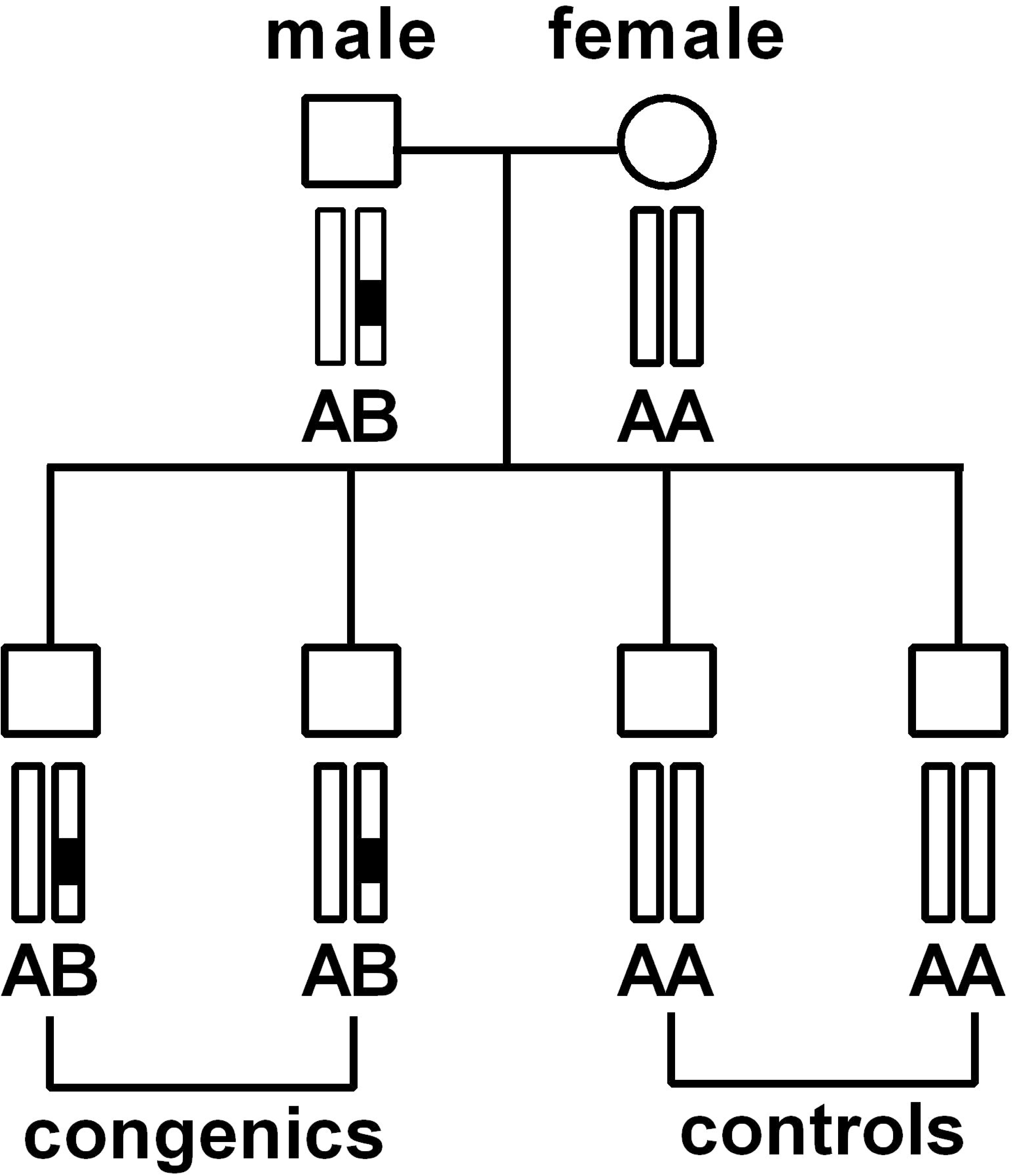
Congenic experimental design to reduce variation due to imprinting and maternal effects. Littermates with one copy of the donor region from the male parent are compared to littermates with the host genotype.

### Body composition

Our primary outcome measure was lean body mass as assessed by dual X-ray absorptiometry (DEXA; PIXImus II densitometer; GE software, version 2.00; Lunar Corp., Madison, WI). We also weighed the body of each mouse to the nearest 0.1 g and measured some but not all congenic mice at 90, 120, 150, and 180 days for lean body mass using magnetic resonance (MR) methods (Bruker mini-spec LF110, Horizontal Whole-Body Composition Rat and Mice Analyzer; Bruker BioSpin Corporation, Billerica, MA). (We obtained this MR instrument in 2012, so mice studied prior to that date have no MR data).

### Genotyping

We assayed genotype of markers on chromosome 2 in a number of ways because the technology changed over the time we bred and studied the mice. For a list of genotyped markers see **S2 Table**. We evaluated simple sequence-length polymorphism markers by polyacrylamide gel electrophoresis after polymerase chain reaction amplification by locus-specific primers [51] in our laboratory. We assayed single nucleotide polymorphisms (SNPs) at three locations: the Genotyping and RNA Analysis Core at the Monell Chemical Senses Center, the Center for Inherited Disease Research (see Electronic Resources) as part of an NIH-funded genotyping supplement and through a commercial vendor (LGC, Beverly, MA; formerly KBiosciences) as a fee-for-service. When assaying variants in the Monell genotyping core, we used primers and allele-specific dye-labeled probes (Life Technologies, Carlsbad, CA). Irrespective of genotyping location and method, controls (blank samples, and genomic DNA from inbred progenitors and their F_1_ hybrids) were included with all assays, and we retested unlikely genotypes as needed. We did not type all mice for all markers, so we imputed missing data by tracing the parental origin of the marker alleles (where applicable) and assumed that no double recombination occurred between markers separated by 26 Mb or less. We cite all genomic base pair positions here relative to GRCm38.

### Data analysis overview

Several goals guided the statistical analysis plan. We wanted to confirm the validity of the varied lean body mass measurement methods, map the *Burly1* locus to the smallest possible physical location, analyze the broader *Burly1* phenotype, and find most or all genes and variants in the *Burly1* region. We describe each goal in turn below.

Before performing parametric statistical analyses, we checked distribution of phenotype for normality within each mapping population using the Lilliefors test and transformed the data as appropriate [52]. All post-hoc tests mentioned below are Fisher’s Least Square Mean tests. For all data analyses, we computed the statistical tests with R (version 3.3.3) and R-studio (version 1.0.136) and graphed the results using either R or GraphPad Prism 6 (version 6.05; GraphPad Software, La Jolla, CA). All data are available for download on Github (http://github.com/DanielleReed/Burly1) and the Center for Open Science (osf.io/yeqjf).

### Validity of measure of lean body composition

We compared the MR, DEXA, and body weight data using Pearson correlation coefficients to determine if these methods gave similar estimates of lean body mass assuming that agreement among methods indicates the validity of each. We focused on male mice for this analysis because we made most of these measures on males. As a further check of the newer MR method, we analyzed the age-related increase in lean body mass expected at 90, 120, 150, and 180 days of age using a repeated-measures ANOVA with these age categories as the repeating measures. (Although we planned to measure mice at these exact days of age, we measured some a few days earlier or later; we grouped mice by age category if they were within 8 days of the target age.) For this analysis, we separated all mice with MR measures into two groups based on genetic background (B6 vs. 129), because mice with a predominantly 129 genetic background are obviously smaller than are those with the B6 background regardless of *Burly1* genotype, and these large differences could mask smaller age effects. We also used a percent score for lean body mass as an outcome with the same analytical approach, computed using day 90 as the baseline: [(lean body mass at age *i* / lean body mass at age 90) × 100]; where *i* = 120, 150, or 180 days.

To map the *Burly1* locus to the physical location, we conducted a general linear model analyses on each of the mapping populations as described below. In addition, we used the common segment analysis method to analyze results obtained from the congenic strain [53].

### Intercross, backcross and congenic analysis

Within each segregating mapping population, we conducted a general linear model analysis with genotype as a fixed factor and body weight as a covariate using a type 1 (sequential) sum of squares. For all populations and for each marker, we calculated (a) the genotype means, (b) the p-value test statistic as the negative base 10 logarithm, and (c) the effect size using Cohen’s *D* [54]. We report these values, including confidence intervals (defined by 2 units of −log_10_ p-value drop), for the peak marker from each mapping population. Statistical thresholds were computed with a Bonferroni correction to an α level of 0.05 for the number of markers (N=122, 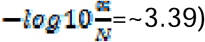. For the first F_2_ population, we used body weight as a proxy measure of lean body mass. We included age as a covariate in any population where age differed by more than a month and if we measured female as well as male mice, we included sex in the model.

### Consomic analysis

Our analysis methods differed between the consomic (B6.129-Chr2) and sub-consomic (129.B6-Chr2) strains. For the consomic strain, we analyzed lean body mass using strain (consomic vs inbred B6) and sex as fixed factors and body weight as a covariate using a type 1 (sequential) sum of squares followed by post hoc tests to determine the significance of strain effects. For the sub-consomic strain, we analyzed the data the same way except using individual genotype at each marker location, i.e., for the sub-consomic mice (129/B6) vs their homozygous littermates (129/129). We calculated the genotype effect size in Cohen’s *D* within each sex for the consomic and sub-consomic strains.

### Congenics and the common segment method

We analyzed the congenic data using the common segment method with the strains listed in **Table 2**. Some strains had very few mice, so we performed two analyses: more broadly, we analyzed all potentially informative congenic strains, defined as strains with at least 3 mice of each genotype group within a congenic strain); more narrowly, we included only the most informative congenic strains, defined as those with at least 38 mice in each of the two possible genotypes per group. These thresholds (3 vs 38 mice) are to some extent arbitrary but are suggested by the actual sample sizes, e.g., some strains have many more mice than others. In addition, we analyzed the effect of the *Burly1* genotype on lean body mass by comparing homozygous congenic mice (strain 2.5; 129/129) with control mice without the 129-derived donor fragment (B6/B6). We conducted this analysis for lean body mass using genotype of marker *rs3666533* as a fixed factor and with body weight and age as covariates using a type 1 (sequential) sum of squares followed by post hoc tests to determine significance of genotype effects using p<0.01 as the significance threshold. We chose this particular marker because it was most strongly associated with lean body mass and we included age as a covariate in this model (**Table 1**).

**Table 1.** Characteristics of the 2,070 mice used in mapping studies

| Population | N | Age range (days) | Tests performed | | | | | Age at MR (days, M±SD) | Period of Study (mm/dd/yr) | | Ref |
| --- | --- | --- | --- | --- | --- | --- | --- | --- | --- | --- | --- |
|  |  |  | BW | DEXA | MR | Metab | OGTT |  | Start | End |  |
| F<sub>2</sub> - First | 397 | 94-354 | √ | x | x | x | x | 195±87 | 09/13/94 | 05/28/96 | [34] |
| F<sub>2</sub> - Second | 113 | 231-270 | √ | √ | x | x | x | 241±9 | 05/28/02 | 07/01/02 | [49] |
| N<sub>2</sub> (F<sub>1</sub> x 129) | 100 | 287-298 | √ | √ | x | x | x | 293±2 | 08/29/05 | 11/23/05 | NA |
| N<sub>2</sub> (F<sub>1</sub> x B6) | 92 | 288-298 | √ | √ | x | x | x | 292±2 | 09/06/05 | 12/12/05 | NA |
| N<sub>3</sub> [(F<sub>1</sub>x129)x129] | 79 | 322-335 | √ | √ | x | x | x | 327±3 | 07/17/06 | 01/17/07 | NA |
| N<sub>3</sub> [(F<sub>1</sub>xB6)xB6] | 49 | 322-335 | √ | √ | x | x | x | 325±3 | 07/17/06 | 01/22/07 | NA |
| 129.B6-Chr2 | 81 | 178-190 | √ | √ | p | x | x | 180±2 | 12/30/09 | 02/22/13 | [41] |
| B6.129-Chr2 | 63 | 175-209 | √ | √ | p | x | x | 187±11 | 11/28/10 | 09/26/14 | [41] |
| Inbred | 49 | 179-207 | √ | √ | p | x | x | 183±5 | 03/27/10 | 01/04/14 | NA |
| Congenic | 1030 | 161-351 | √ | √ | √ | p | p | 188±28 | 11/16/11 | 04/05/16 | NA |

Population, type of mapping resource; N, number of mice; Age range for the last test of mice. BW, body weight; DEXA, dual X-ray absorptiometry; MR, magnetic resonance; Metab, metabolism measured using the LabMaster equipment; OGTT, oral glucose tolerance test; Age at MR, final age point only (mean ± standard deviation (SD)); most of mice underwent MR at 180 days, but some were a few days older or younger); 'Start', start date for breeding and 'End', end date for breeding by month/day/year; Ref, reference that describes breeding of the mapping population. √, measured in all mice; p, partial (measured in some but not all mice); x, not measured in that population; NA, not applicable.

**Table 2.** Characteristics of congenic strains

| Strain | N | Age at DEXA (days) |  | Tests Performed |  |  |
| --- | --- | --- | --- | --- | --- | --- |
| | | Range | Mean±SD | MR | Metab | OGTT |
| 1 | 218 | 165-200 | 180±3 | x | x | x |
| 1.1 | 2 | 181-181 | 181±0 | x | x | x |
| 1.10 | 8 | 179-180 | 179±0 | p | x | x |
| 1.11 | 4 | 180-180 | 180±0 | √ | x | x |
| 1.11.1 | 77 | 179-182 | 180±0 | p | x | x |
| 1.12 | 8 | 179-181 | 180±0 | p | x | x |
| 1.13 | 12 | 180-181 | 180±0 | p | x | x |
| 1.14 | 1 | 180-180 | 180±0 | √ | x | x |
| 1.2 | 14 | 180-181 | 180±0 | p | x | x |
| 1.3 | 3 | 180-180 | 180±0 | x | x | x |
| 1.4 | 9 | 180-181 | 180±0 | p | x | x |
| 1.5 | 10 | 180-180 | 180±0 | √ | x | x |
| 1.6 | 1 | 180-180 | 180±0 | √ | x | x |
| 1.7 | 17 | 180-183 | 180±0 | p | x | x |
| 1.7.1 | 29 | 179-224 | 187±16 | √ | x | x |
| 1.8 | 11 | 180-181 | 180±0 | p | x | x |
| 1.9 | 8 | 179-180 | 179±0 | p | x | x |
| 2 | 164 | 177-192 | 180±2 | x | x | x |
| 2.1 | 16 | 180-250 | 204±32 | p | x | x |
| 2.2 | 2 | 181-181 | 181±0 | √ | x | x |
| 2.3 | 27 | 180-350 | 195±34 | p | x | x |
| 2.4 | 171 | 179-183 | 180±0 | p | x | x |
| 2.5 | 195 | 161-351 | 216±50 | p | p | p |
| 2.6 | 10 | 180-181 | 180±0 | √ | x | x |
| 2.7 | 10 | 180-180 | 180±0 | √ | x | x |

N, number of mice; SD, standard deviation (an SD of 0 indicates less than one day); DEXA, dual X-ray absorptiometry; MR, magnetic resonance at four time points (90, 120, 150, and 180 days); Metab, metabolism measures using the LabMaster equipment; OGTT, oral glucose tolerance test. √, measured in all mice; p, partial (measured in some but not all mice); x, not measured in that strain. Owing to husbandry errors, we made DEXA measures outside of our normal procedures that are not reflected in the table totals: strain 2.2, n=2 measured at 120 days old; strain 2, n=1 measured at 100 days old and these 3 mice were not included in this table; homozygous strain 2.5, n=36 measured at 8 months of age and n=15 at 161-163 days old; n=7 for homozygous congenic with donor (129/129) and n=44 their littermates without donor (B6/B6). For the metabolism measures, we measured 52 mice in total, n=30 with the B6/129 genotype and n=20 with the B6/B6. For the OGTT, we measured 47 mice in total, n=25 with the B6/129 genotype and n=22 with the B6/B6.

### Age

To determine how early in adult life we could detect the effects of *Burly1,* we applied a repeated-measure ANOVA using lean body mass both in grams and as a percentage of baseline for mice from the relevant congenic strains. (By ‘relevant’ we mean the congenic strains with donor regions that overlapped with the *Burly1* locus, defined as genotype variation at marker *rs3666533*.) In total, 21 *Burly1* congenic strains were included in this analysis. We excluded four congenic strains because of incomplete MR data (strains 1, 1.1, 1.3, and 2).

### Metabolism

We examined whether mice that differed in lean body mass as function of *Burly1* genotype also differed in food intake, activity, or metabolism. Using specialized equipment (TSE LabMaster, version 5.0.6; TSE Systems, Inc., Chesterfield, MO, USA), we measured mice from congenic strain 2.5 and the control group from the littermates without 129 donor fragment (B6/B6). We chose this congenic strain because it contained the smallest donor region that contained the *Burly1* locus. To measure these traits, we trained mice for several days in cages that mimicked the experimental cages to ensure they learned to eat and drink appropriately. When we transferred the mice to the experimental cages, we quantified food intake and water intake corrected for lean body mass, physical activity in three dimensions including rearing as well as walking, and increases in carbon dioxide production and decreases in oxygen consumption. We used these values to compute heat produced per hour for each mouse, correcting for lean body mass rather than total body weight [55], and we expressed all data as the mean of four 24-hr data acquisition cycles. To analyze these data, we used t-tests to compare genotype groups of marker *rs3666533* (129/B6 vs B6/B6 in the congenics)

### Oral glucose tolerance test (OGTT)

To determine if the *Burly1* genotype affects oral glucose tolerance, we measured 22 heterozygous congenic mice (strain 2.5; 129/B6) and a control group of their 25 homozygous (B6/B6) littermates. We deprived mice of food for at least 4 hours but not longer than 18 hours and gavaged them with 0.2 g/ml glucose solution for a final dose of 2 g/kg mouse body weight. We collected tail blood samples twelve times (baseline, 5, 15, 20, 25, 30, 45, 60, 75, 90, 105, and 120 minutes post-gavage) and measured blood glucose concentrations (mg/dL) using an Accu-Chek Avia Plus meter (Roche Diagnostics, Indianapolis, IN, USA). Missing data points were replaced with average values within each genotype group at the same time point. We analyzed the data using a repeated one-way ANOVA with a type 1 (sequential) sum of squares followed by post hoc tests.

### Evaluating genes and variants in the Burly1 region

Drawing on the genomic coordinates suggested by the congenic results from the common segment analysis method, we found all previously annotated genes within the *Burly1* region using an online database [56]. We used another online database [57] to find genomic variants among inbred mouse strains related to the B6 and 129 stains studied here [58, 59]. We formatted this information using an online tool [60] and we identified those regulatory and coding variants with the potential to cause functional changes [61]. In addition, we identified human genes and their variants associated with body mass that are located in the region of conserved synteny with the mouse *Burly1* region by searching an online catalog of human genome-wide association results [62] using the key word ‘lean body mass’ as well as the less specific term ‘body mass index’.

## Results

### Overview

In **Table 1** we list the number of mice studied per mapping population, their age range, and which of the three lean body mass methods we used to measure this trait. In **Table 2** we list the individual congenic strains, number of mice studied per strain and their ages at DEXA analysis as well as which additional measures we made on these mice. The wide range in the number of mice bred for each congenic strain owes to breeding difficulties and practical constraints on the size of our animal colony. Here we show details of every congenic strain we bred even those which were potentially uninformative because of the sample size. However, we eliminated the data from three mice: two were pregnant (owing to husbandry errors), and one had a large kidney tumor.

### Normality

We assess whether lean body mass data were distributed normally within each mapping population. Significant deviations from normality were present in the first F_2_ and the pooled congenic population. For the first F_2_, no transformation was effective at normalizing the distribution of the data and because no method achieved the desired result, we report the analysis of the untransformed data after confirming that the results were similar using all of the transformations attempted (**S3 Table**).

### Validity

Lean body mass was validly measured by both DEXA and MR. We draw this conclusion because the three measures—body weight, and lean body mass measured by DEXA and by MR— while not identical, were highly correlated (**Figure 2**; r-values, 0.62-0.95; p<0.0001). The exact r-statistics varied depending on the mapping population. **S1 Figure** shows all body weight, DEXA, and MR correlation data; **S4 Table** provides all correlation test statistics by mapping population. For the primary end-point measure, we used the DEXA data, which provided a more complete analysis because we did not have MR data for all mice.

**Figure 2.**
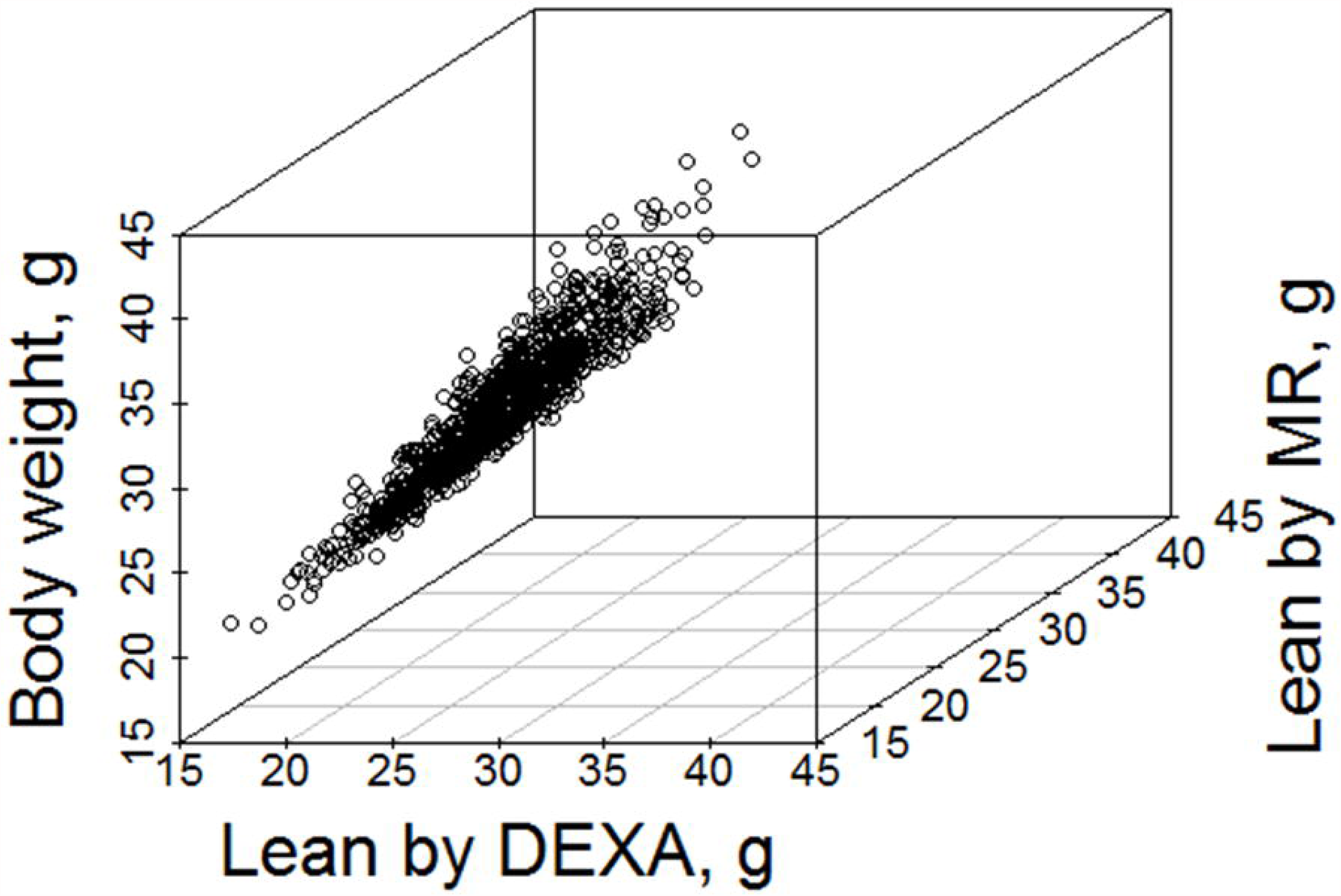
3D scatter plot of body weight and lean body mass measured by DEXA and MR in male mice. These measures are highly correlated (r-values of 0.62-0.95, p<0.0001) within mapping populations (**S1 and S2 Figure**).

In addition, as we expected, mouse lean body mass increased over time until ∼150 days of age and was stable thereafter (**S2a and b Figure**). We also noted that, independently of the *Burly1* locus, mice with a 129 genetic background differed in the pace of lean body mass growth from those with a B6 background, especially between 120 and 150 days of age. Mice with the 129 background were still adding lean body mass during this period, whereas those with the B6 background plateaued (**S2c Figure**).

### Mapping

For the intercross, backcross, and congenic mice, we show the association test statistics and the QTL locations, with confidence intervals, for each mouse mapping population (**Figure 3a-e**), as well as the genotype mean of lean body mass (**Figure 3f-j**) and the effect sizes for the peak linked marker (**Figure 3k-o**). We found a single common genomic region (around 152 Mb) responsible for lean body mass in all mapping populations (**Figure 3a-e**). There is a consistent direction of allelic effect with the B6 allele increasing lean body mass relative to the 129 allele (**Figure 3f-j**). In some cases the effect sizes attributable to genotype were large, over 0.5 in backcrosses with the B6 background and among the congenic strains (**Figure 3n, o**). The reciprocal backcrosses have a similar *Burly1* effect, but the effect is larger in mice with the B6 rather than 129 genetic background (**Figure 3c, d, h, i, m, n**). The *Burly1* effect was also similar in both reciprocal consomic/sub-consomic strains. The introgression of chromosome 2 from the 129 strain (B6.129-Chr2) reduced lean body weight relative to the inbred parent strain mice (**Figure 4a**). These results further confirm the observation that the B6-derived allele increases lean body mass (**Figure 3f-j**) and we also learn that it does so in both male and female mice. In fact, the effect size is larger in females than in males (**Figure 4b**).We observed similar genotype and sex effects from the reciprocal sub-consomic mice (**Figure 4c-e**).

**Figure 3.**
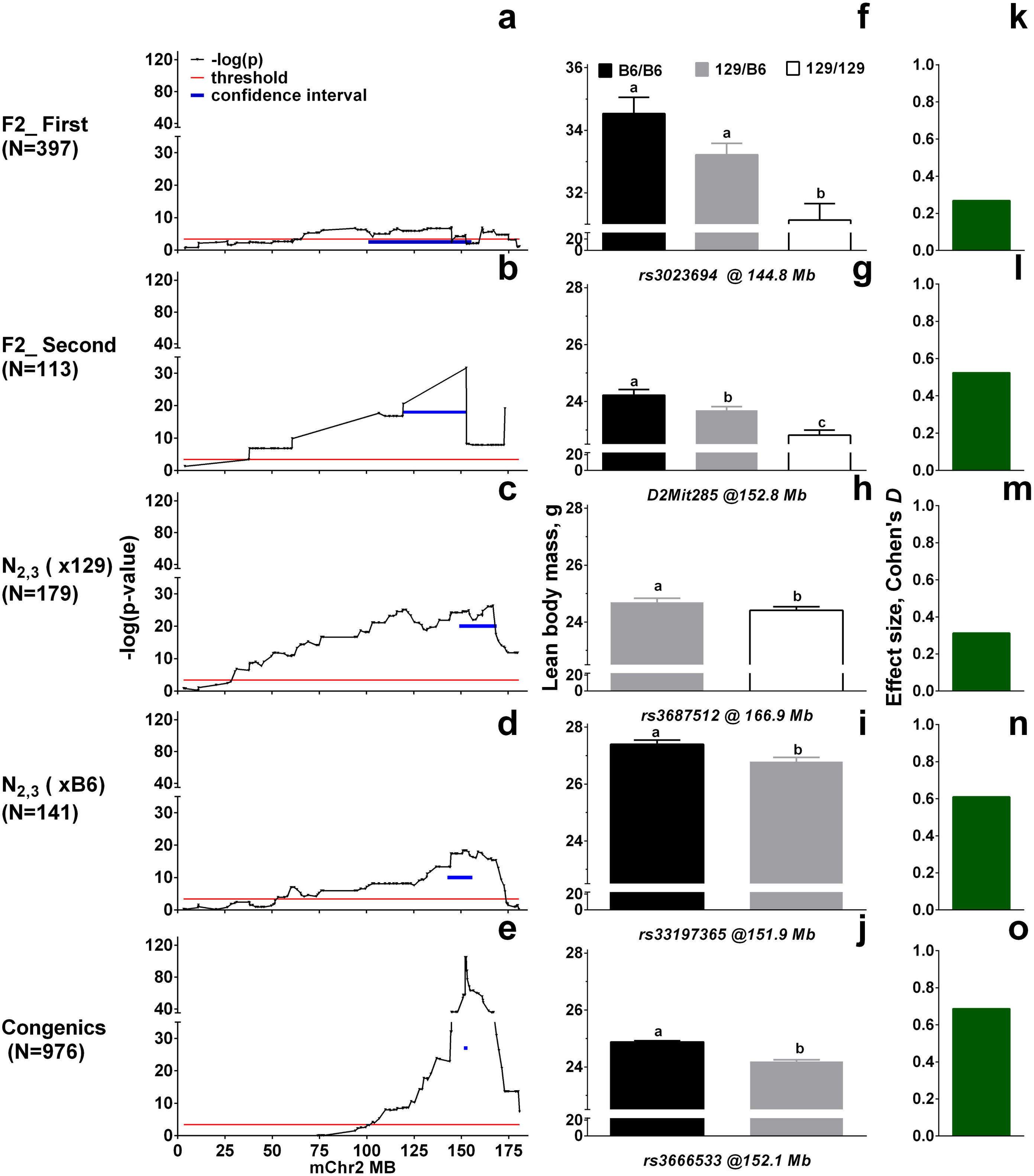
The genomic location of mouse QTL *Burly1* identified in multiple mapping populations. (**a-e**) Association test statistics and QTL locations, with confidence intervals, for each mouse mapping population. The x-axis is the location of the markers in Mb on mouse chromosome 2 (mChr2); the y-axis is the −log of the statistic test by each marker genotype. The blue bars indicate the confidence intervals of the QTLs that were supported by 2 units of −log_10_ p-value drop. The red horizontal line shows a Bonferroni-corrected statistical threshold. (**f-j**) Mean and standard error of lean body mass of mice grouped by peak marker genotype. For the first F_2_ population the results are for body weight not lean body mass. Letters a, b, and c show significant differences between genotypes (p<0.00001, post hoc tests, general linear model). (**k-o**) Effect sizes of lean body mass at the peak marker for each mapping population. For the two F_2_ populations, the effect size was calculated in Cohen’s *D* calculated using least square means of genotypes of B6/B6 vs. 129/B6, which allows us to compare it with the congenic mice with 129 donor fragment onto the B6 host inbred strain.

**Figure 4.**
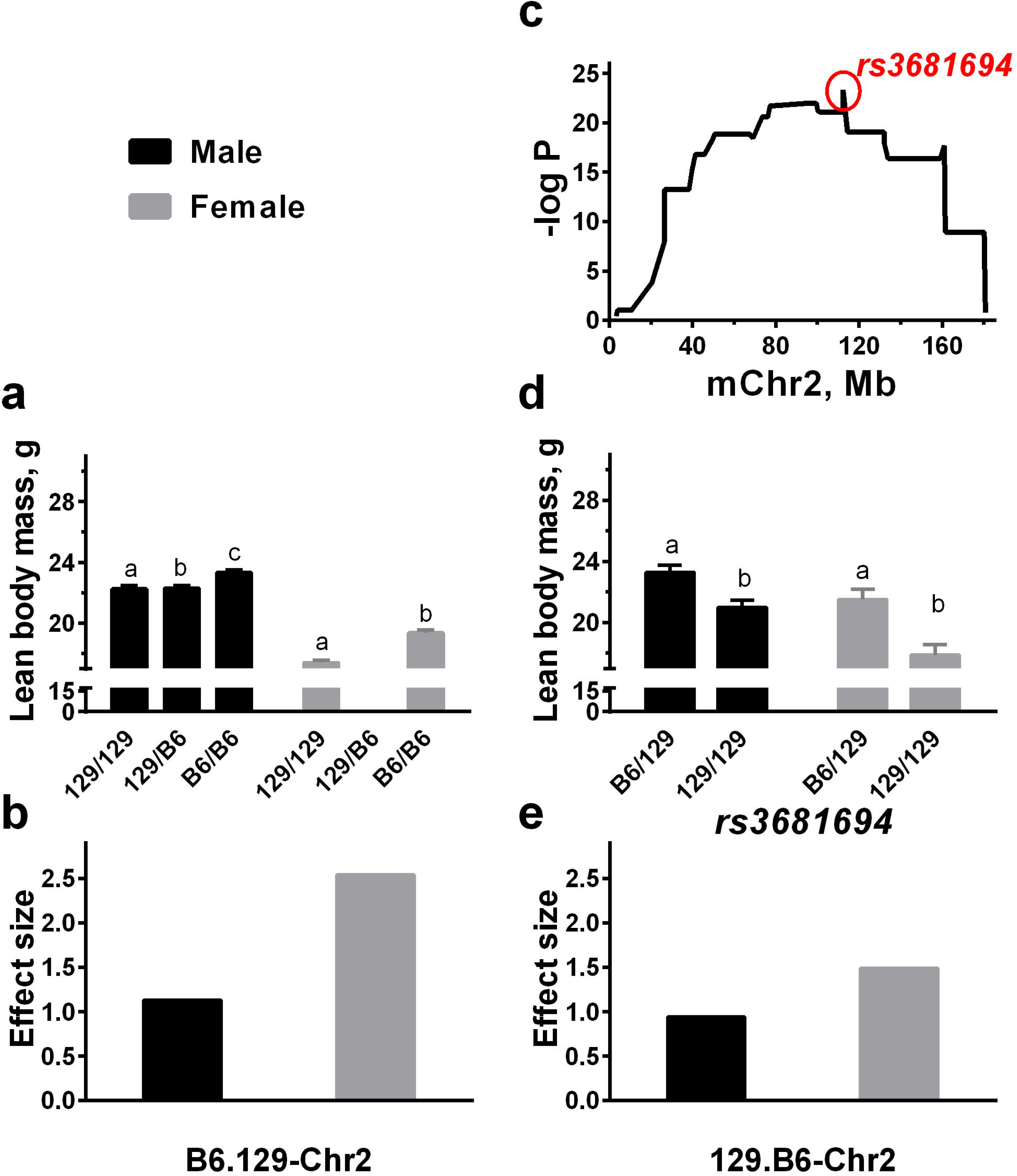
The *Burly1* locus has consistent effects in the reciprocal consomic and sub-consomic strains. (**a, b**). B6.129-Chr2 consomic strain: homozygous, heterozygous (male only) consomic, and inbred host strain (C57BL/6ByJ). Means ± SEM lean body mass (a) and effect size in Cohen’s *D* (b) were computed using B6/B6 vs. 129/129. (c) Mapping of the sub-consomic 129.B6-Chr2 mice (N_6_, N_7_, and N_10_) created as part of the consomic process shows broad linkage peaks. The x-axis is the location of the markers in Mb on mouse chromosome 2 (mChr2); the y-axis shows results of a general linear model using genotype and sex as fixed factors and body weight as a covariate for lean body mass as the −log p-value (black line). The strongest associated marker is *rs3681694* (red). (**d, e**) 129.B6-Chr2 sub-consomic strain: backcross mice grouped by *rs3681694* genotype: heterozygous mice with one copy of the B6 allele (129/B6) vs. homologous littermates without a B6 allele (129/129). Data are means ± SEM (d) and the effect size in Cohen’s D (e) computed using B6/129 vs. 129/129. Letters a, b, c show significance at p<0.05 by post hoc testing.

### Congenics

We bred 1,030 congenic mice from 25 congenic strains with donor regions of varying lengths and breakpoint locations (**S5 Table**), which we confirmed by genotyping each congenic mouse. Not all strains were equally informative for the common segment method because of sample size so did two analysis, one narrow and one broad. (For a list of strains with inclusion by analysis method, see **S6 Table**). We conclude based on both the narrow and broad approaches that there is a 0.8-Mb region of chromosome 2 that contains the *Burly1* locus (151.9-152.7 Mb; **Figure 5a**, **S5 Table**). We draw this conclusion because using the general linear model, there is distinct and highly significant peak at that location (**Figure 5a**), and because from the common segment approach, this region is shared among the strains with the *Burly1* genotype effect and not shared with strains without this effect (**Figure 5b**). We show the results by individual congenic strains in **Figure 5c** and all post-hoc tests (including those for the broad analysis below) in **S7 Table**. **Figure 5d** shows all known noncoding RNA, protein-coding genes, pseudogenes, and processed transcripts within this region obtained from the Ensemble Mouse Genome Brower. The broader results also pointed to the same region (**S3 Figure**). Likewise, the broader analysis of the common segment method pointed to the same physical location on the chromosome with the same direction of allelic effect (**Figure 5c**, **S3c Figure**).

**Figure 5.**
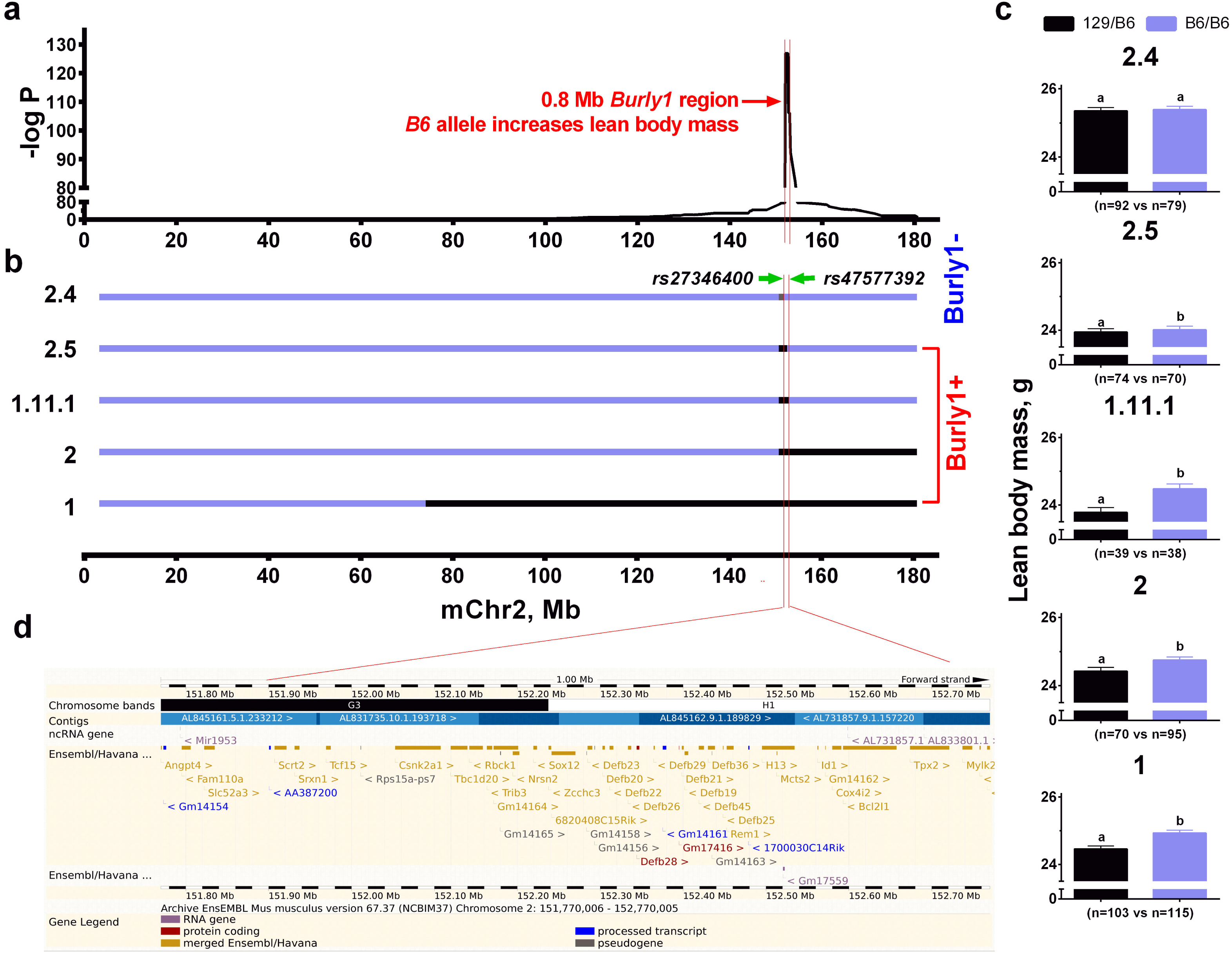
The *Burly1* locus region isolated by comparing the five most informative congenic strains. (a) Average lean body weight compared using a general linear model with body weight as a covariate among all congenic mice grouped by genotype at each marker. The x-axis is marker positions in Mb on chromosome 2 (mChr2); y-axis, −log_10_-transformed p-values. (**b**) Donor region of each congenic strain: black bar, strain retained the *Burly1* locus; gray bar: strain did not retain the locus. Blue indicates the region contributed by the host strain. We determined whether congenic strains (shown at left) retained the *Burly1* locus (i.e., were ‘positive’) by comparing within each strain the average lean body mass of littermates with and without the donor fragment. (**c**) Strain comparisons with sample size (n) of each genotype within each strain. *p<0.05/18=0.002778 except for strain 2.5, p=0.013416; ns: p>0.05. (d) *Burly1*-positive strains share a common region (red line; 0.8 Mb from rs33197365at 151.9 Mb to rs3700604 at 152.7 Mb) that the *Burly1*-negative strains do not share. The allele effect direction matches that from the consomic mice, with the B6 strain allele increasing the trait. We show noncoding RNA genes, protein-coding genes, pseudogenes, and processed transcripts within the 0.8 Mb *Burly1* region which we obtained from the Ensemble Mouse Genome Brower.

Analysis of the homozygous congenic strain and control mice showed consistent results for the *Burly 1* location and direction of effect (strain 2.5, **S5 Table**). Mice homozygous for the 129 allele had 1.5 grams less lean body mass compared with mice with the B6/B6 genotype [F(1,47)=10.477, p=0.002] (129/129 vs B6/B6). We used this particular mapping resource because of its small donor region to demonstrate that the *Burly1* effect is on lean and not on fat mass [F(1,47)=0.064, p=0.801; **Figure 6**]. This result is consistent with an analysis of fat mass rather than lean body mass using the general linear approach with all informative congenic strains (**S4 Figure**).

**Figure 6.**
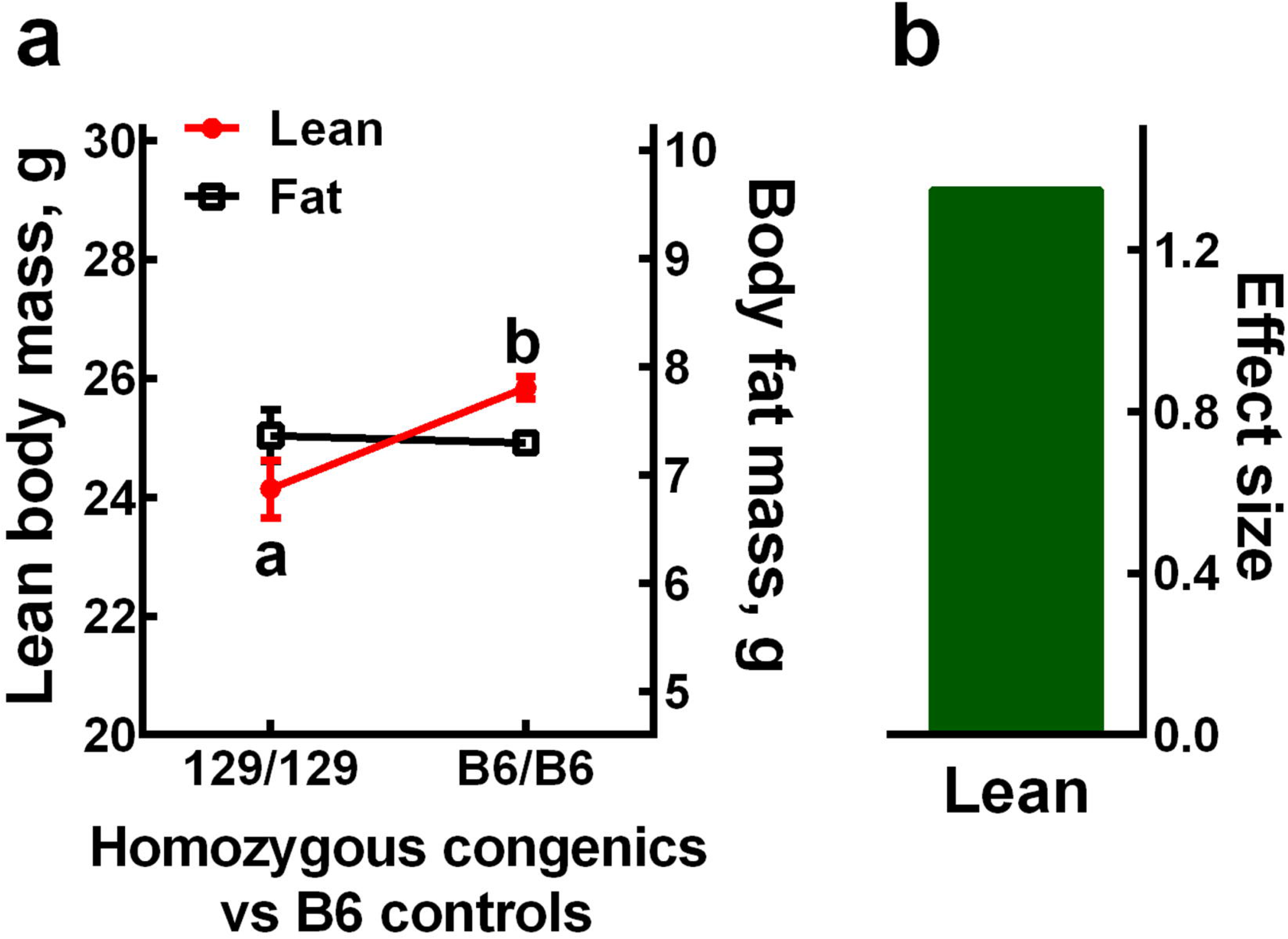
The phenotype effect of the lean-body-mass-specific *Burly1* locus is confirmed in homozygous congenic strain 2.5. (**a**) Mean ± SEM of lean body mass (red circles) and body fat mass (black squares) for homozygous congenic mice vs. mice descended from their littermates. We found a significant genotype effect on lean body mass (p<0.0001, post hoc tests) but not body fat mass. (**b**) Effect size in Cohen’s *D* for lean body mass.

### Age by genotype

The *Burly1* locus affected lean body mass at every time point we measured, 90, 120, 150, and 180 days of age, with the B6 allele consistently increasing the phenotype (p<0.001, repeated-measures ANOVA with post hoc comparison) and a similar effect size (Cohen’s *D*) at all age points (0.40, 0.56, 0.45, and 0.45, respectively; **Figure 7**). We analyzed percent growth to understand the rate of lean body mass increase, but this measure did not differ by genotype [F(1,303)=2.6, p=0.11, repeated-measures ANOVA]. These results suggest mice with the *Burly1* B6 allele gain more lean body mass than those with the relevant 129 allele in early life (prior to our first measurement at 90 days old) and that this pattern persists through the window of time measured here.

**Figure 7.**
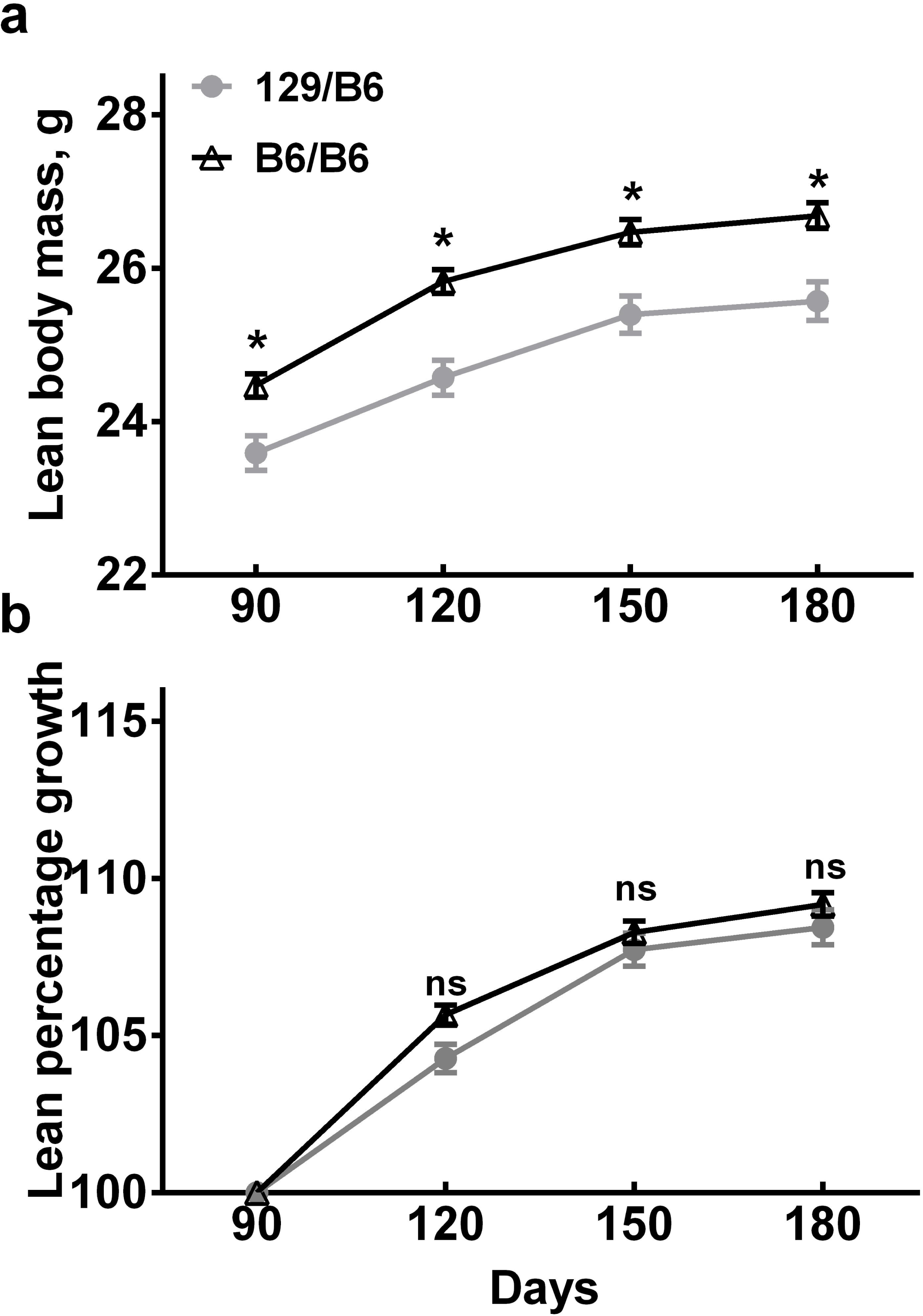
The *Burly1* genotype effect on lean body mass in male mice at ages 90, 120, 150, and 180 days. Sample sizes for genotypes of marker *rs3666533* are n=209 for B6/B6 and n=96 for 129/B6. *Burly1* significantly affects lean body mass, with B6 allele consistently increasing the trait (**a**); *p<0.001, repeated-measures ANOVA with post hoc tests), but we observed no genotype effect on monthly lean body percentage growth at all age points (**b**).

To check whether the *Burly1* locus affected fat mass, we reanalyzed congenic strains using fat mass as the outcome measure. Consistent with results from the homozygous congenic strain, these additional results show that the *Burly1* locus is independent of fat mass (**Figure 6**), although there is a nearby fat mass QTL (**S4 Figure**). The congenic strain 2.5 (**S5 Table**) retains the *Buly1* locus but no loci that affect fat mass (**Figure 6**, **S4 Figure**).

### Metabolism

We studied the metabolism of 52 congenic mice and littermates with opposing genotypes (stain 2.5; **Table 2**). We display the results in **Figure 8**. Mice with the 129-derived allele consumed less the oxygen [ml/kg lean, hr; t(1,49)=2.143, p=0.037], and produced less the heat per kilogram lean body weight [kcal/kg lean, hr; t(1,50)=2.179, p=0.034] and the carbon dioxide [ml/kg lean, hr; t(1,50) =2.032, p=0.047] than did their littermates without 129- drived allele. However, there were no significant *Burly1* genotype effect on food [g/kg lean, 24 hr; t(1,49)=1.073, p=0.289] and water [ml/kg lean, 24 hr; t(1,49)=0.625, p=0.535], respiratory exchange ratio [ t(1,50)=0.743, p=0.461] or activity [infrared beam breaks/hr; t(1,50)=0.278, p=0.782]. Genotype also did not account for differences among mice in plasma glucose concentration at any time point after they were gavaged with glucose [F(1, 45)=0.93, p=0.347; **S5 Figure**].

**Figure 8.**
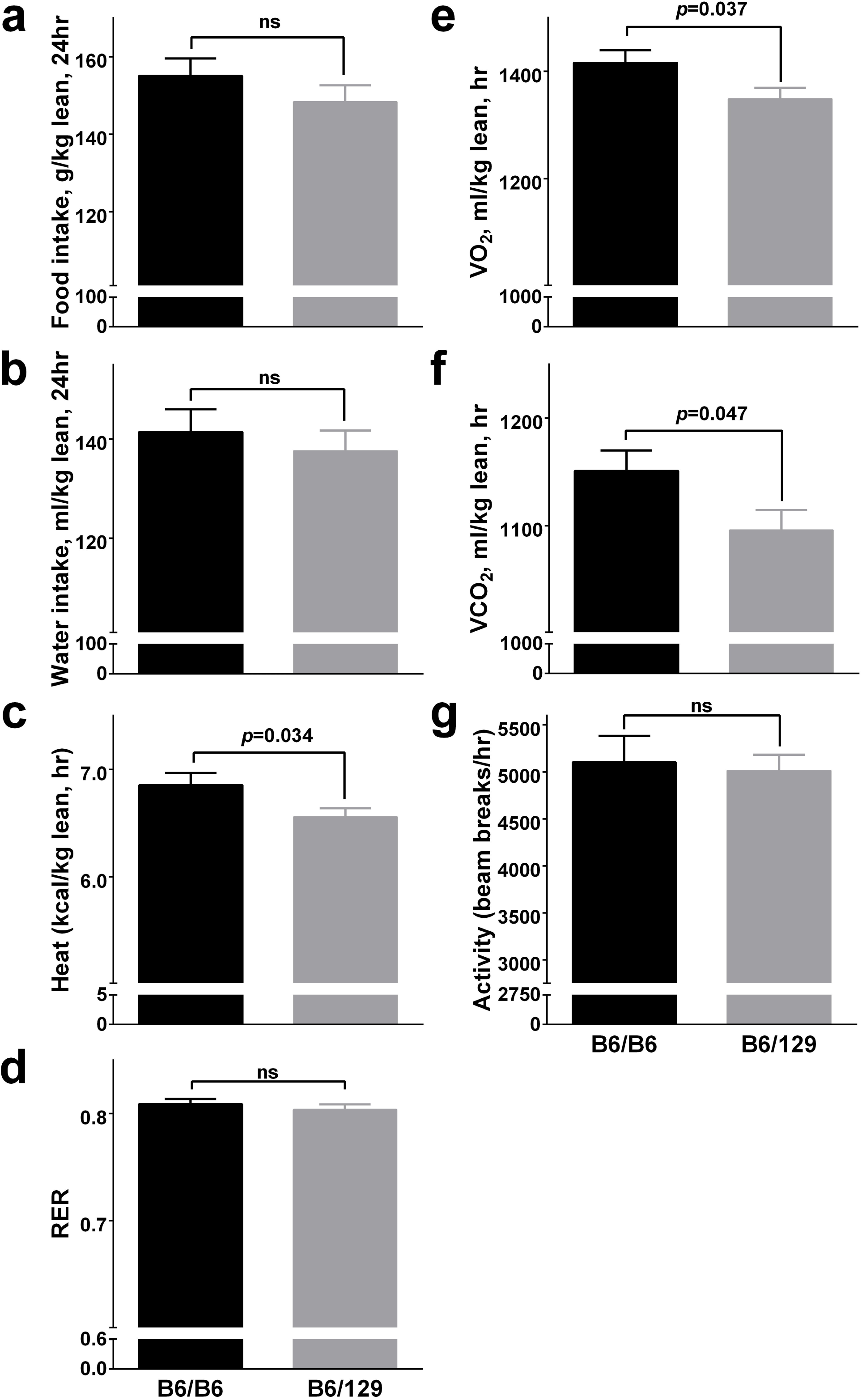
Metabolic assessments in the *Burly1* congenic strain 2.5 and control mice: food intake (a), water intake (b), heat production (c), respiratory exchange ratio (d), oxygen consumption (VO_2_; e), carbon dioxide production (VCO_2_; f), and activity over the entire 24-hr light/dark cycle (g). Data are mean ± SEM.

### Candidate genes

Using the coordinates suggested by the congenic strain mapping results, we surveyed the Mouse Genome Database [3] for other body weight and lean body mass QTLs on chromosome 2. The eight previously reported QTLs (*Gnf1, Wg2d, Bwq9, BWq5, Pwbwq1, Pwgrq1, Pwgrq2, and Pwbwq5*) had an overlapping confidence interval with the *Burly1* QTL reported here (**S8 Table**). We extracted 1949 variants between the B6 and 129 strains (**S6 Figure**). Of these variants, the *in silico* analysis predicted that 7% change some aspect of mRNA regulation (**S9 Table**) and 2.5% affect nonsense-mediated mRNA decay (**S10 Table**). There are 41 protein-coding genes [56] (**S11 Table**), seven of which contain missense variants (**S12 Table**). To the best of our knowledge, none of these variants is within a gene previously studied for its affect in the development or regulation of lean body mass, as determined by searching a publicly available database of experimental studies using the gene symbols as search terms [63]. Using the genomic coordinates of the *Burly1* region (Chr2:152009210- 152613619), we identified the homologous region of the human chromosome (chr20: 142056-759014) and compared recent human genome-wide association results for lean body mass or body mass index to determine if the regions contained variants in common genes, but we found none.

## Discussion

### Overview

Positional cloning of body composition loci during the genomic era was initially quick, most notably the identification of several obesity genes, including leptin [64], the leptin receptor [65], tubby [66, 67], and agouti [68]; however, progress identifying QTLs with smaller effect sizes or with complex architecture has been slower. For example, we found in an attempt to narrow the causal allele for an adiposity QTL (*Adip20*) that it decomposed into several linked QTLs with intra-chromosomal epistasis [69]. We were surprised that the *Burly1* phenotype mapped to a single region because adiposity and related traits have many interacting and sub-QTLs on mouse chromosome 2. However, the *Burly1* locus has an easily detected and consistent effect on body composition that maps to a small region on chromosome 2, with the B6 allele increasing lean mass compared with the 129 allele.

### Control of nongenetic factors

Lean body mass is affected by many factors such as age, sex, and diet [70] so we took steps to reduce these sources of variation whenever possible. For the strains studied here, we measured male mice almost exclusively and took care to compare mice that were close in age whenever practically possible. In retrospect, our choice to study adult mice from 3 - 6 months of age was fortuitous because the *Burly1* genotype effects are large and consistent during this time window. Therefore, the choice of age to study was not a limitation; however, studying mostly male mice was a limitation, and this choice reduced our ability to generalize our findings to female mice [71], especially because the two sexes differ in similar studies [72]. However, we did examine a few female mice, and learned that the *Burly1* phenotype of females was similar and perhaps even surpassed that of males.

Like age and sex, diet is another source of variation. To control these effects, we fed mice food ordered from the same manufacturer with the same catalog number for the duration of this project. However, we acknowledge that chow diets change over time because of their natural constituents (e.g., grains) [73]. Therefore, the vicissitudes of diet composition could contribute to nongenetic sources of variation, especially for mice studied years apart.

### Challenges in assessing body composition loci on mouse chromosome 2

Mouse chromosome 2 has posed special challenges for the genetic study of body composition because of its QTL density [37, 38, 40, 72, 74-78] and the interdependence of body composition traits. QTL density was a consideration when we initially chose among the available fine-mapping approaches [79, 80]. We adopted the congenic approach because this method seemed most suitable for isolating very closely linked QTLs [81], and because of our experience with it [82]. The tight interdependence of body composition traits was also a challenge. Lean and fat mass are correlated in mice [83], as they are in humans [84, 85], so we chose methods that measure lean and fat weight separately. One additional challenge was that, while we measured lean body mass and the weight of many organs, we did not measure muscle weight directly. The lack of a direct measure was unfortunate, especially considering the presence of a nearby QTL for muscle weight [86].

### Biological basis of breeding problems

We attempted to create and study reciprocal consomic strains to compare the effect of the *Burly1* allele on two different genetic backgrounds. This is an important goal for the study of body composition because many nonspecific genetic effects reduce body size. Therefore, we wanted to determine not only whether the 129-derived allele reduces body size but also whether the B6-derived allele increases it. However, our breeding plan for these reciprocal strains was only partially successful, because we found it nearly impossible to breed consomic mice with a B6 donor chromosome 2 on a 129 genetic background. These breeding difficulties were unsurprising because prior studies show that the agouti region on chromosome 2 interacts with other loci to reduce reproductive performance [87, 88]. However, we did successfully produce heterozygous mice with a donor sub-chromosome 2; from those mice, we learned that the B6-derived allele from the *Burly1* region increased lean body mass. Thus, this sub-consomic strain, while imperfect, was informative and further confirmed that *Burly1* has specific effects that, depending on the allele, increase or decrease lean body weight.

### Mouse human homology and measurement of body composition

Investigators who have conducted human genome-wide association studies of lean body mass, measured using methods similar to those used here in mice, report no associations to the homologous *Burly1* region [89-94]. However, this may be due to low power of human studies to detect genes with smaller effect sizes because, relative to studies of body mass index, far fewer human subjects have been measured for lean body weight. Again, drawing on body mass index as an example, many additional loci are uncovered as sample size increases [95]. Other explanations for this lack of human-mouse agreement would be that the causal gene has few or no functional variants in humans or may not have the same function between mice and humans [96]. Genetic studies in mice can point to functional roles of genes, and their value is due in part to this knowledge even in the absence of comparable human variation [97].

### Candidate genes

Within the 0.8-Mb *Burly1* region there are at least seven protein-coding genes (*Angpt4, Fam110a, Slc52a3, Zcchc3, 6820408C15Rik, Defb25, Rem1*) with missense variants between two mouse strains closely related to the parental strains we studied here [98]. To the best of our knowledge, investigators have not reported a role for any of these genes in lean body weight, and there have been only a few functional studies of any type. However, there are clues about how a few of these genes might affect lean body weight. Perhaps most compelling, the protein product of the *Angpt4* gene is a secreted growth factor that promotes the growth of blood vessels. Inborn differences in the function or abundance of this growth factor may affect the amount of lean tissue, although there is no direct evidence for this hypothesis that we are aware of except that the degree of *Angpt4* gene methylation differs by body weight in humans [99]. The *Slc52a3* gene codes for a protein that transports riboflavin [100], and treatment with riboflavin in humans with mutations of this gene improves the strength of their muscles by acting at the motor neuron [101]. It is possible that improving or reducing the nerve-muscle junction will increase or decrease overall lean body mass. The *Rem1* gene codes for a GTP-binding protein that inhibits a particular type of voltage-dependent calcium channel in muscle [102]. Following the logic applied above, reducing muscle function might change lean body mass. The protein produced from the *Fam110a* gene is a member of a small family of proteins that form part of the centrosome when cells divide, and while it is not specific to muscle cells, it might affect the pace of cell division and perhaps final cell number [103].

From the characterization of the congenic mice, we learned that *Burly1* genotype has no effect on food and water intake when expressed per unit of lean body mass, which is similar to the results of our previous studies of the progenitor strains [104, 105]. However, the *Burly1* congenic mice with a 129-derived allele produce significantly less heat per kilogram of lean body weight than did controls. This current result is consistent with a prior observation that 129 vs. B6 strain variation affects heat-generating mitochondria within brown adipocytes in muscle [106].

### Future work

Investigators have made progress is understanding gene function by making mice with knockout alleles and studying the effects on multiple traits [107]. While a useful method in general, null alleles can have nonspecific effects on body size, and we have observed that when gene knockout is not immediately lethal, at least a third of genes when nullified affect body weight, usually reducing it [22, 23]. Therefore, this method gives little insight into the effects of naturally occurring variation, which typically has more nuanced effects on protein or other functions. Making changes to the genome is now much easier with the new gene editing technology, which has been used in many species, including yeast [108-110], zebrafish [111], fruit flies [112], nematodes [113], plants [114], mice [115], monkeys, and even human embryos [116, 117]. This method works well when we know one or only a few target alleles and can produce and compare them, but it is of limited help when there are hundreds of potential causal variants to evaluate, as is the case here. Thus, we need better ways to prioritize genes and variants from positional cloning studies in mice and genetic association studies of humans to understand how specific variants affect traits like lean body mass.

## Acknowledgments

We gratefully acknowledge the assistance with animal breeding of Rebecca James, Liang-Dar (Daniel) Hwang, Zakiyyah Smith, Matt Kirkey, Amy Colihan, and Laurie Pippett. We also acknowledge Richard Copeland and the consistent high-quality assistance of the animal care staff at the Monell Chemical Senses Center, and thank them for their service. Michael G. Tordoff and Gary K. Beauchamp commented on a draft of the manuscript.

---

**S1 Figure.** Correlations among three measures, body weight and lean body mass measured by DEXA and MR, in male mice of each mapping population. (**a**) Comparison of MR and DEXA results. (**b**) Comparison of body weight and lean body mass measured by MR (red data points) or DEXA (green data points). Despite the small sample size of two populations (B6.129-Chr2 and C57BL/6ByJ), the three measures are highly correlated (r-values = 0.62-0.95, p<0.00001). B6.129-Burly1 refers to all congenic mice from **Table 2**.

**S2 Figure.** Monthly lean body mass increases in male mice from two populations (mice with 129 background, n=13; and mice with B6 background, n=319). (a, b) Monthly lean body mass increased significantly (*p<0.05) from 90 to 180 days for both strains. There is no significant difference (p=0.84) in lean body mass between 150 and 180 days for the 129 strain. (c) Monthly percentage of lean body mass, fitted to a sigmoid growth model, increased robustly before 150 days of age and then stabilized. The differences in percentages between mice with 129 background and B6 background are significant (*p<0.05) at 150 and 180 days.

**S3 Figure.** The *Burly1* locus region was isolated by comparing the 18 informative congenic strains. (**a**) Average lean body weight compared using a general linear model with body weight as a covariate, among all congenic mice grouped by genotype at each marker. The x-axis shows marker positions in Mb on chromosome 2 (mChr2); y-axis, −log_10_-transformed p-values. (**b**) We determined which congenic strains retained the *Burly1* locus (i.e., were ‘positive’) by comparing within each strain (shown at left) the average lean body weights of littermates with and without the donor fragment. Black bars, donor region retained the *Burly1* locus; gray bars, donor region did not retain the locus; blue bars, region contributed by the host strain. For the three strains labeled with the red $, there was no reliable genotype effect on lean body mass. *Burly1*-positive strains share a common region (red lines; 0.8 Mb from *rs33197365* at 151.9 Mb to *rs3700604* at 152.7 Mb) that *Burly1*-negative strains do not share. (**c**) Comparison of allele effect across strains. The allele effect direction matches that from the consomic mice, with the B6 allele increasing the trait.

**S4 Figure.** The *Burly1* is a lean-body-mass-specific locus that has no effect on body fat mass. (**a**) Average body fat weight compared using a general linear model with body weight as a covariate, among all congenic mice within each congenic strain grouped by genotype at each marker. The x-axis shows marker positions in Mb on chromosome 2 (mChr2); y-axis, −log_10_-transformed p-values. The blue bar shows the confidence interval of the fat locus, defined by a drop of 2 units of −log p-value. (**b**) The 0.8 Mb *Burly1* region defined in the congenic strains, which is out of the fat locus region (blue bar in a). (**c**) A significant genotype effect on body fat mass was found only in strains 1 and 2 (red stars in b) that retain two largest 129-derived donor fragments, and no genotype effect was found in the other 16 strains. Thus, the location of the fat locus differs from the *Burly1* region. *p<0.05; post hoc tests.

**S5 Figure.** No *Burly1* genotype response to oral glucose tolerance tests in mice and their littermates’ control from congenic strain 2.5. Mean ± SEM of genotype was reported and the significance of genotype effect was evaluated by post hoc tests using p=0.05 as a significance level.

**S6 Figure.** Statistical category of variants within the *Burly1* region based on their predicted variant effect.

